# Overt attention towards appetitive cues enhances their subjective value, independent of orbitofrontal cortex activity

**DOI:** 10.1101/648931

**Authors:** Vincent B. McGinty

## Abstract

Neural representations of value underlie many behaviors that are crucial for survival. Previously, we found that value representations in primate orbitofrontal cortex (OFC) are modulated by attention, specifically, by overt shifts of gaze towards or away from reward-associated visual cues (McGinty et al., 2016). Here, we investigate the influence of overt attention on behavior, by asking how gaze shifts correlate with reward anticipatory responses, and whether activity in OFC mediates this correlation. Macaque monkeys viewed Pavlovian-conditioned appetitive cues on a visual display, while the fraction of time they spent looking towards or away from the cues was measured using an eye tracker. Also measured during cue presentation were the monkeys’ reward anticipation, indicated by conditioned licking responses (CRs), and single neuron activity in OFC. In general, gaze allocation predicted subsequent licking responses: the longer the monkeys spent looking at a cue at a given time point in a trial, the more likely they were to produce an anticipatory CR later in that trial, as if the subjective value of the cue were increased. To address neural mechanisms, mediation analysis measured the extent to which the gaze-CR correlation could be statistically explained by the concurrently recorded firing of single OFC neurons. The resulting mediation effects were indistinguishable from chance. Therefore, while overt attention may increase the subjective value of reward-associated cues (as revealed by anticipatory behaviors), the underlying mechanism remains unknown, as does the functional significance of gaze-driven modulation of OFC value signals.

## INTRODUCTION

Neural value representations underlie many of the behaviors we rely on to survive, from simple appetitive and defensive reflexes to complex economic decisions. Several recent studies have shown that value representations in the prefrontal cortex can be influenced by how attention is allocated among visual objects of different value. This includes overt shifts of attention (gaze) performed during natural free viewing (McGinty et al., 2016; Hunt et al., 2018), as well as covert shifts of attention performed in the absence of saccadic eye movements (Xie et al., 2018). Allocation of gaze also influences economic choice behavior, with increased gaze time on a given item making it more likely to be chosen over the alternatives (Krajbich et al., 2010; Towal et al., 2013; Vaidya and Fellows, 2015; Gidlöf et al., 2017). A natural hypothesis emerging from these studies is that attention, by modulating neural value signals, may influence a wide range of value-driven behaviors. To test this hypothesis, we build upon our recent report of gaze-modulated value signals in the primate orbitofrontal cortex during appetitive Pavlovian conditioning (McGinty et al., 2016). Whereas the prior report considered only the neural effects of gaze, here we address both the neural and behavioral effects, and ask whether the neural effects are sufficient to explain behavior.

Pavlovian conditioning is a form of learning in which otherwise neutral cues acquire motivational significance (value) after being paired with pleasant or aversive outcomes, so that presentation of the cues alone can elicit conditioned responses (CRs). These responses are usually stereotyped, reflexive behaviors performed in direct anticipation of the outcome (e.g. salivation in anticipation of food), and can vary according to the size, probability, frequency, or desirability of the predicted outcome. Our central hypothesis is that overt attention influences CRs performed in anticipation of reward, and that attentional modulation of OFC is the mechanism underlying this influence.

To test this hypothesis, we simultaneously measured Pavlovian CRs, eye movements, and OFC neural activity, in an appetitive conditioning task as described previously (McGinty et al., 2016). We then asked whether trial-by-trial variability in gaze allocation towards the cues corresponded to variability in CR magnitude, and whether this correlation could be statistically explained (mediated) by the firing of single OFC neurons. Although the effect varied according to subject and trial condition, in general we observed a positive correlation between gaze and CRs: The longer the monkeys spend looking at a Pavlovian cue in a given trial, the more likely they are to perform a CR later in that trial, suggesting that gaze allocation influences the in-the-moment subjective value of the cue. With respect to the role of the OFC, we found no evidence that OFC activity could explain the correlation between gaze and CRs, suggesting that some neural substrate outside of the OFC must mediate the influence of gaze on reward anticipation.

## METHODS

### Overview

Macaque monkeys performed an appetitive Pavlovian conditioning task (Figure 1A) while three variables were measured simultaneously: allocation of gaze (overt attention), reward anticipation, and value representations in OFC (Figure 2). Gaze location was measured relative to the location of reward-predictive Pavlovian cues, and gaze allocation was quantified as the fraction of time the monkeys spent looking at the cues. Reward anticipation was defined as the conditioned licking responses (CRs) that monkeys performed in the moments leading up to reward delivery, and was quantified as the fraction of time that a CR response was detected. OFC value representations were measured on the basis of single- and multi-unit neural activity (see Analysis below).

**FIGURE 1.**
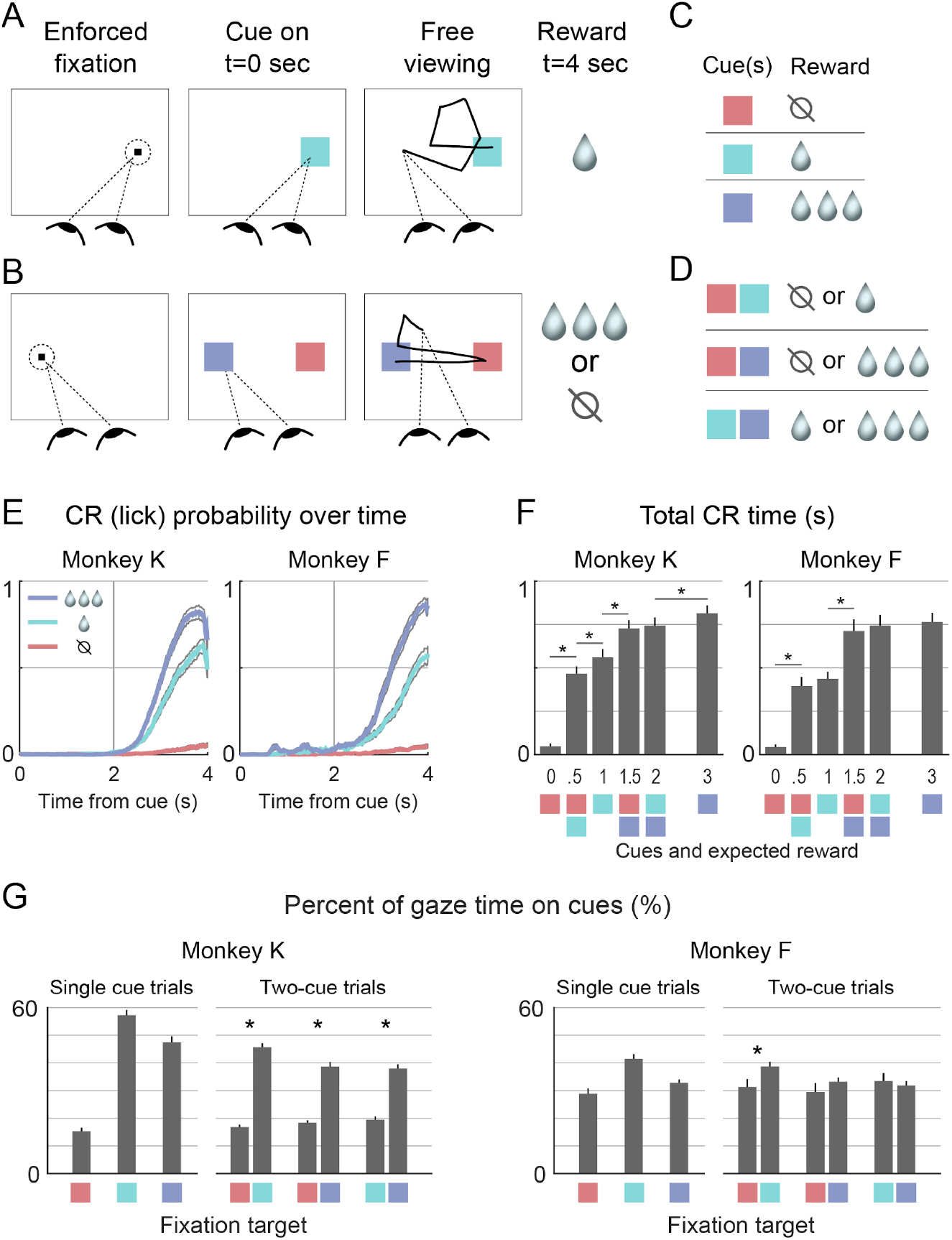
(A-D) Monkeys were trained to associate visual cues with juice reward. Single cues were followed by certain reward (A,C), and cues shown in pairs were followed by probabilistic rewards (B,D). Anticipatory conditioned licking responses (CRs) and allocation of gaze onto the cues were tracked in every trial. (E) CRs in single-cue trials. The y-axis gives the probability of a CR being detected at a given time point. Colored and gray lines give the mean and 95% confidence intervals, respectively. (F) CRs in all trial types. The y-axis gives the total time that a CR was detected during cue presentation. The six bars in each graph correspond to the six trial types, with colored squares below the x-axis to indicate the cue configuration, and numerals to indicate the mean juice value of the cue(s). (G) Allocation of gaze onto cues in all trial types. The y-axis gives the percent of time that the gaze was on (within 3 degrees of) the cue center. Single-cue trial types are indicated by the three single colored squares, corresponding to the cue value. Two-cue trial types are indicated by adjacent colored squares, corresponding to the cue values shown in that trial type; in these trials, gaze allocation was tallied separately for each cue, hence two bars for each two-cue trial type. Note that gaze allocation was dependent on cue type, but was not always a monotonic function of value. Data in E-G reflect 25 sessions for Monkey K and 28 for Monkey F. Bars show means across sessions, and whiskers show SEM. * indicates p<0.05 (corrected) in a comparison between immediately adjacent bars.

**Figure 2:**
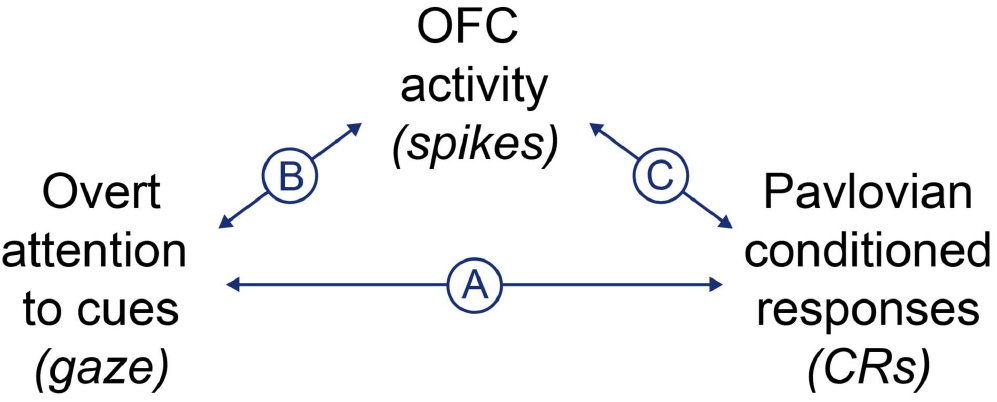
Overview of major analyses: (A) trial-by-trial correlation between gaze allocation and CRs. (B) modulation of OFC activity by gaze, and (C) trial-by-trial correlation between OFC activity and CRs. We also perform a mediation analysis, which quantifies the degree to which the gaze-CR correlation (A) can be statistically explained by the combined effect of gaze on OFC activity and the correlation between OFC activity and CRs (B and C).

The analyses had two main objectives. The first was to determine whether gaze allocation and reward anticipation were correlated with one another on a trial-by-trial basis. The second was to determine whether this correlation could be explained, in a statistical sense, by the activity of OFC neurons. To satisfy the first objective, we computed the correlation between the time spent looking at Pavlovian cues and the duration of CRs across trials (Figure 2A). Importantly, the relatively long trial duration (4 seconds) allowed the correlation to be assessed across different time points in the trial; thus, we were able to determine whether looking at a cue early in the trial was correlated with CRs later in the trial, i.e. whether gaze allocation could predict subsequent reward anticipation. The results are shown in Figure 3 and 4.

**Figure 3.**
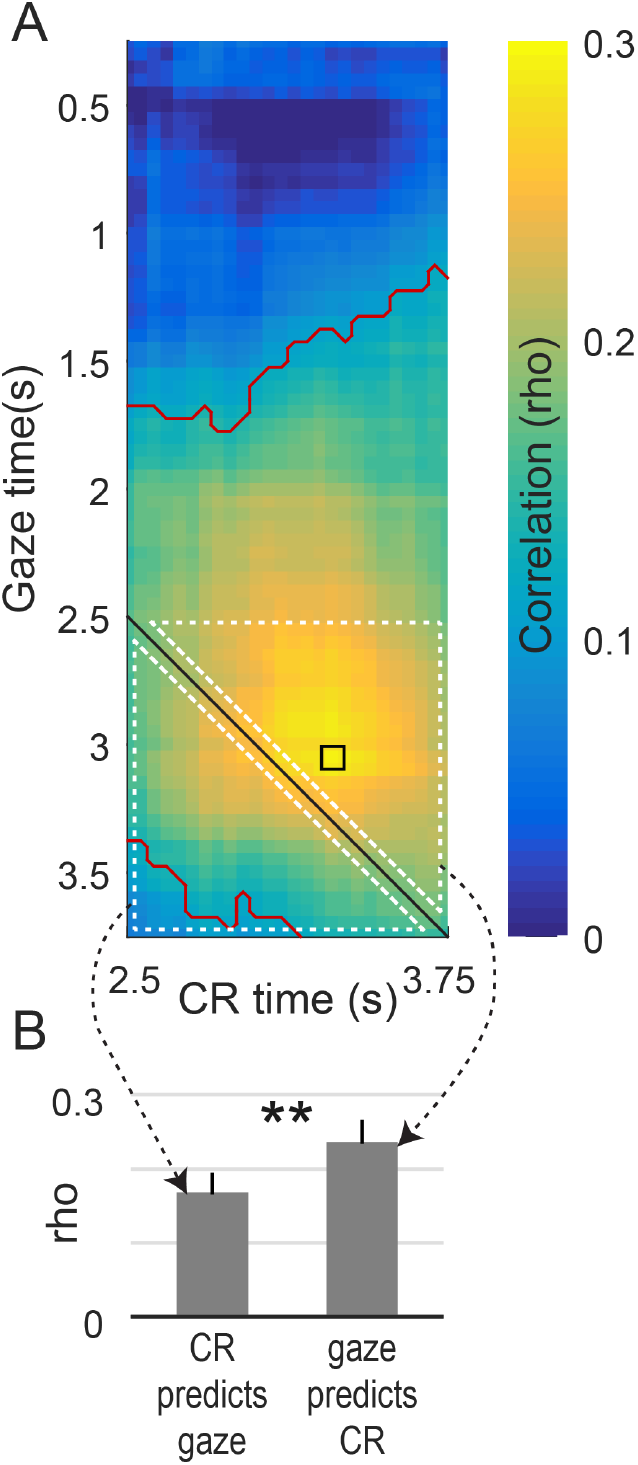
Gaze allocation predicts CRs, in example data from a single trial type in one monkey. (A) The heatmap shows the trial-by-trial correlation between time spent looking at a cue and time spent performing an anticipatory CR, assessed across different time points in the trial. Warmer colors indicates that greater gaze allocation was associated with more CRs. The x- and y-axes give the times at which the CR and attentional data were observed, respectively, and equivalent time points are given by the black diagonal. The small black square shows the peak correlation. Red contours indicate mean significantly above zero (mean of 25 sessions, p<0.001 with cluster correction at p<0.01). White dotted triangles indicate time points averaged together to produce the bar graphs in panel B. (B) Bar heights give the average correlation within the white-outlined pixels above and below the black diagonal in A. Whiskers give the SEM, and ** indicates significant difference between the bars (p<0.01, corrected, n=25 sessions, paired t-test).

**Figure 4:**
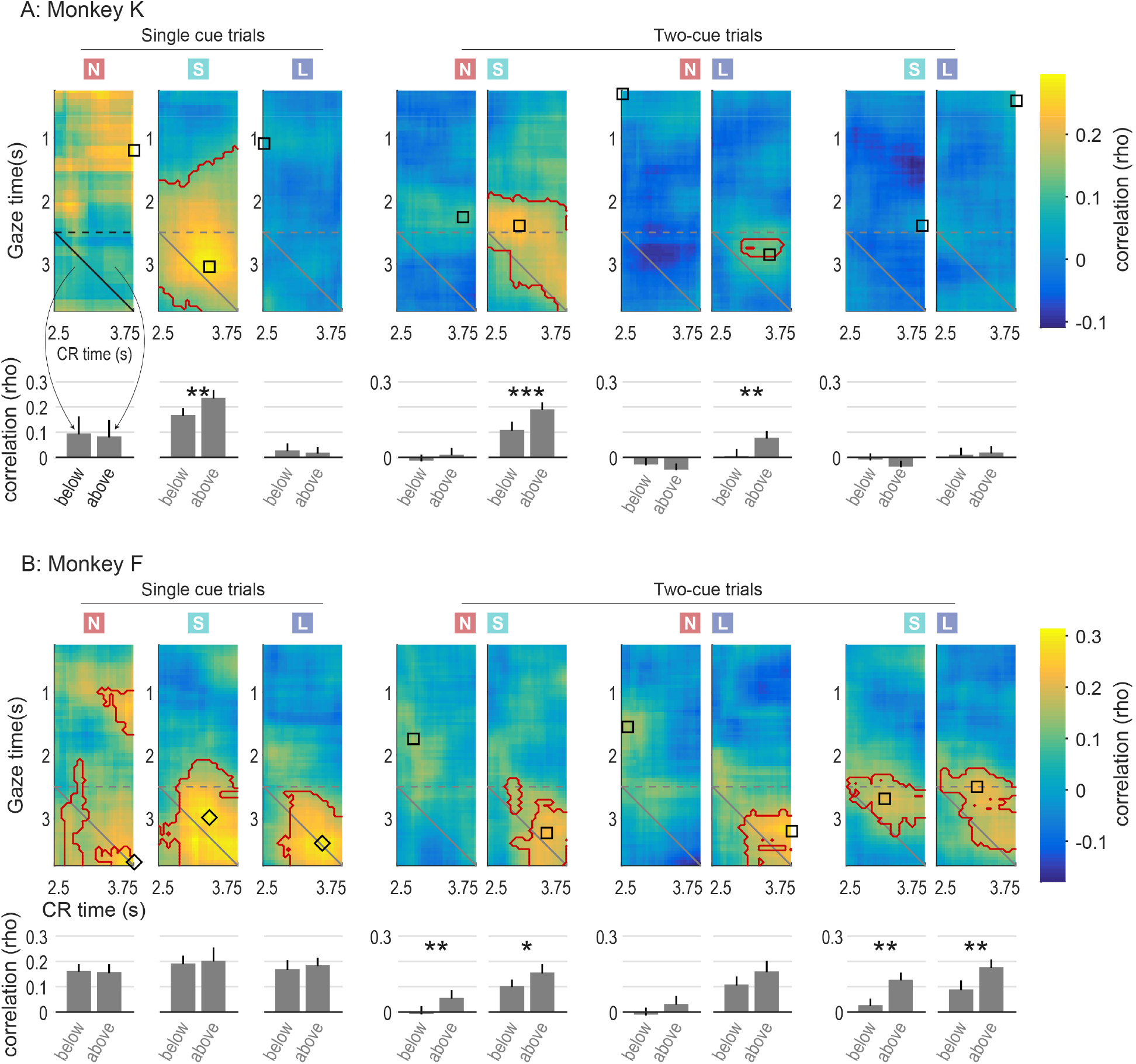
Gaze-CR correlation in all trial types for both monkeys. The colored squares at top indicate whether the gaze data pertain to ‘none’, ‘small’, or ‘large’ value cues (red N, green S, and blue L, respectively). Note that in two-cue trial types, gaze is assessed separately for each cue, hence there are two conditions for each two-cue trial type. Heatmap colors and contours follow the same conventions as Figure 3A, and bar graphs below each heatmap follow the same conventions as Figure 3B. The black or diamond on each map indicates the time point used in subsequent analyses of OFC spiking activity (see main text and Figure 8). Squares show the point of highest gaze-CR correlation in the map, and diamonds show the highest point above the main diagonal (see Methods). *, **, and *** indicate significance between adjacent bars, at p<0.05, p<0.01, and p<0.001, corrected. The data from Figure 3 are reproduced in the first row, second column (single ‘small’ cue trials in Monkey K).

To satisfy the second objective required testing two correlational relationships: between gaze and OFC activity (Figure 2B), and between OFC activity and CRs (Figure 2C). The relationship between gaze and OFC activity was assessed using linear models, as in our prior work (McGinty et al., 2016) (results in Figures 5 and 6). To quantify the relationship between OFC firing and CRs, we used a modified form of Spearman’s correlation coefficient (results in Figures 7 and 8). Finally, to quantify the degree to which OFC firing could statistically explain the gaze-CR correlation, we performed a mediation analysis (results in main text). These last two analyses were also performed separately within each trial type.

**Figure 5:**
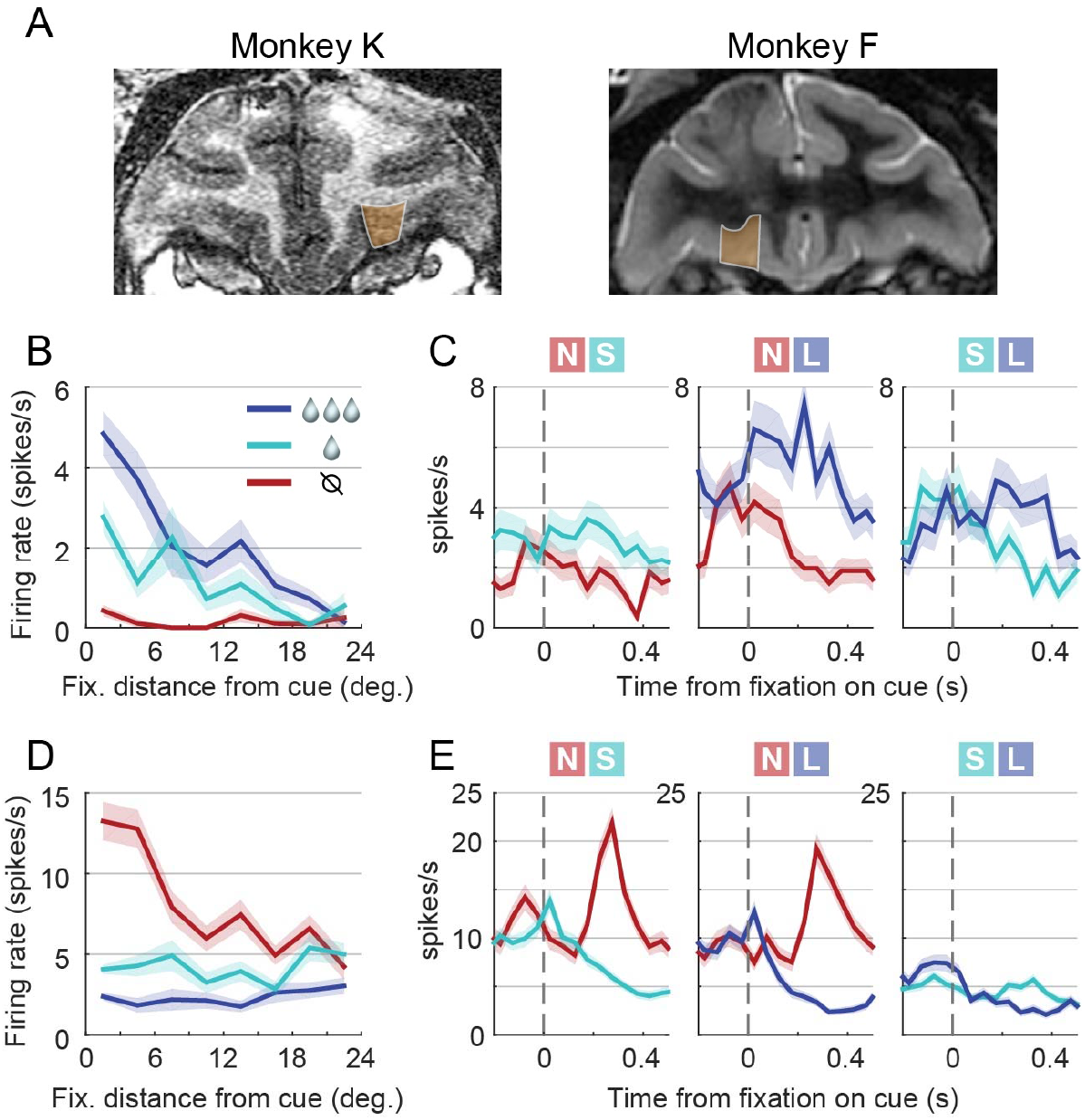
Value signals in OFC neurons were modulated by shifts of gaze towards or away from the cues. (A) Coronal MRI sections from the two subjects; orange shading shows the recorded areas. (B) Single example cell. In single-cue trials, firing increased as a function of cue value (colors) and decreased as a function distance of fixation from the cue (x-axis). In addition, the effect of value was maximal for fixations on or near the cue, constituting an interaction between the value and distance effects. (C) Same cell as in B. During two-cue trials, firing was greater following fixations onto the higher of the two cue values shown. Each of the three panels shows firing time-locked to fixation onset at t=0; the trial type (cues shown) is at top, and the line color indicates which of the two cues was fixated. Lines give the means, and shading gives the SEM. (D, E) A second example cell, showing effects of value and attention (including interaction effects) on firing in single cue trials (D) and two-cue trials (E). Unlike the cell in B and C, firing decreases as a function of single cue value, or of the fixated cue value in two-cue trials.

Most analyses were performed separately for each trial type; for example, trials with a single ‘no reward’ cue were analyzed as a group, separately from trials with a single ‘small reward’ cue, and separately from trials with both a ‘no’ and ‘small’ reward cue shown simultaneously, etc. This was done because the correlation between gaze and CRs (the key behavioral outcome in this study) differed markedly between trial types, as we illustrate below. However, to identify OFC cells modulated by gaze, all single-cue trial types were analyzed together, and all two-cue trial types were analyzed together. This was done in order to assess the encoding of value, which could only be done by comparing firing across trials with differing cue value (i.e. different trial types).

### Subjects and apparatus

All procedures were performed in accordance with the NIH Guide for the Care and Use of Laboratory Animals, and were approved by the Animal Care and Use Committee of Stanford University, which was where the data were collected. The subjects were two adult male rhesus monkeys designated K and F, weighing 13.5-15.0kg. Using aseptic techniques, they were implanted with an MR-compatible head holder, and subsequently with a recording chamber (Crist Instruments, Hagerstown, MD); a craniotomy was also performed to allow access to the OFC. Data were collected while the monkeys were head-restrained and seated ~57 cm from a fronto-parallel CRT monitor displaying the task stimuli. The stimuli were square color patches (3.2 degrees per side) and were mutually isoluminant. Horizontal and vertical eye position was recorded at 400Hz. A tube for fluid rewards was placed outside the mouth, and to retrieve an available reward the monkeys had to touch their tongue to the end of the tube during delivery. Both monkeys quickly learned to do so, and typically consumed all of the juice delivered on every trial. Monkeys typically performed anticipatory conditioned licking responses prior to reward delivery, and these were quantified according to the fraction of time that a response was detected in a given epoch (see below).

Task flow and stimulus presentation were controlled using the REX software suite (Laboratory of Sensorimotor Research, National Eye Institute) and dedicated graphics display hardware (Cambridge Research Systems). Neural signals were measured from single tungsten electrodes (FHC Inc., Bowdoin, ME) placed at the target locations using a motorized drive (NAN Instruments, Nazareth, Israel). Neural activity, eye position, and task event data were acquired and stored using a Plexon MAP system (Plexon, Inc., Dallas, TX).

### Behavioral Task

The task was identical to that used in McGinty et al (2016), with the exception that on some trials, two cues were shown simultaneously. See figure 1A-D for an illustration, and below for details. The monkeys were trained to associate three different color cues with three juice rewards in approximate ratios of 3:1:0. These are referred to as “large”, “small”, and “none”, or as “L”, “S”, and “N”, and they are indicated in the figures by the colors blue, turquoise, and red. Juice volumes were constant within a session, but varied slightly across sessions to compensate for changes in the monkeys’ fluid sensitivity during the study. A session was defined as the behavioral and neural data collected on a single day; more than one cell was typically recorded in each session. Only sessions with concurrently recorded neural data were used.

Trials began with a fixation point (FP) appearing 5 degrees to the left or right of the screen center. After the monkey fixated on this point for 1-1.5 seconds, either one or two cues were shown, at which point the monkey was free to move his eyes. Eye position was monitored, but had no consequence for trial outcome. Reward was delivered 4 seconds after cue onset, depending on which cue or cues were shown (see below). The cue(s) was extinguished at 4.3s after cue onset, after which there was a 2-4s inter-trial interval, followed by the illumination of the FP on the next trial.

Trials had either a single cue, or two different cues shown simultaneously. In single cue trials (Figure 1A), one randomly chosen cue appeared at the location of the FP, and the volume of reward delivered at the end was determined by the color of the cue (Figure 1C). In two-cue trials, one randomly selected cue appeared at the FP location (5 degrees left or right of center), and a different randomly selected cue appeared 5 degrees from center in the opposite direction of the FP location (Figure 1B). At the end of the trial, one of the two reward volumes was randomly chosen to be delivered (Figure 1D). For example, the trial illustrated in Figure 1B has a “large” and “none” cue, indicating a 50% probability of a large reward, and a 50% probability of no reward. Single-cue and two-cue trials types were presented in equal proportions, randomly interleaved within a session.

New cue colors were selected for every session by randomly sampling equidistant points on a color wheel. Each session therefore began with a learning phase, which was completed before data collection. During learning, single cue trials were presented until the monkey’s CR during cue presentation (the 4 seconds prior to reward) became proportional to the reward size: large > small > none, with the CR for “none” trials being absent or negligible. Learning was considered complete when the CR durations were significantly different (rank sum test, p<0.01 uncorrected) over the previous 60-100 learning trials. Learning phase data were not used in any analysis in this paper. In a prior report from this data set (McGinty et al., 2016), we examined the effects of cue-reward “reversals” on some OFC cells. Those cells are also used here; however, for a given cell we use only the pre- or post-reversal data (never both) according to whichever segment of the data had more trials. In other words, in this report the cue-reward associations were static, with no reversals.

### Conditioned responses and quantification of reward anticipation

Monkeys typically performed conditioned licking responses (CRs) during the 4 second cue display period in anticipation of reward delivery. CRs were quantified by detecting the presence/absence of contact between the tongue and juice delivery tube. This was done by connecting the input lead of a single channel amplifier (A-M Systems, 400Hz sampling) to the fluid reservoir, and the ground lead to the seat of the monkey’s chair. Tongue contact with the juice tube abruptly reduced the amplitude of ambient noise on the channel, and setting an appropriate noise threshold effectively binarized the signal into epochs of contact/no contact. The CR-vs.-time plots in Figure 1E show the proportion of trials in which contact was present at a given time point. Total contact time throughout the trial was averaged across trials of a particular type to produce Figure 1F. For the analyses in Figures 3, 4, 7, and 8, CR data were first segmented into overlapping bins, each 500ms in duration, with 50ms increments between bin centers. The first bin was centered at 2500ms after cue onset, because CRs were nearly always absent until that time (Figure 1E). The last bin was centered at 3750ms after cue onset, for a total of 26 bins. CRs were then quantified by finding the fraction of time within each 500ms bin that contact was detected.

### Eye tracking and quantification of gaze

Gaze was unrestricted during the 4s cue display period. Horizontal and vertical eye position were recorded at 400Hz using a non-invasive optical system in Monkey K (Eyelink, SR Research), and scleral search coil system in Monkey F (C-N-C Engineering). These different eye tracking methods yield similar data (Kimmel et al., 2012)

Gaze location was quantified in relation to the cue or cues. For Figures 3, 4, 7, and 8, gaze data were segmented into overlapping bins (500ms each, 50ms increments between bin centers), over the 4 second cue presentation period, yielding a total of 71 bins with the first centered at 250ms after cue onset and the last centered at 3750ms. Gaze allocation for each bin was quantified as the fraction of time (out of 500ms) that gaze was within 3 degrees of the center of a cue (‘on’ the cue). In two-cue trial types, gaze allocation was tallied for each cue individually.

In Figures 5 & 6, OFC neural activity was analyzed with respect to gaze location. Here, it was necessary to time-lock neural data to gaze behavior, to create a temporal reference point for peri-event time histograms (PETHs, as in Figure 5C,D), and to account for the known temporal lag between visual events and OFC activity (Thorpe et al., 1983; Wallis and Miller, 2003; Padoa-Schioppa and Assad, 2006). Therefore, eye position data were segmented into fixation and saccade epochs (Engbert and Kliegl, 2003; Kimmel et al., 2012), and the fixation onsets were used as the temporal reference point for spiking data, as in our prior work (McGinty et al., 2016). To assess neural data with respect to fixation location in single-cue trials, fixations were quantified according to the distance of gaze from the cue center (e.g. Figure 5B,D), consistent with our prior report. In two-cue trial types, fixations away from the cues were infrequent, and so neural analyses only used data from ‘on-cue’ fixations (within 3 degrees of the cue center).

**Figure 6:**
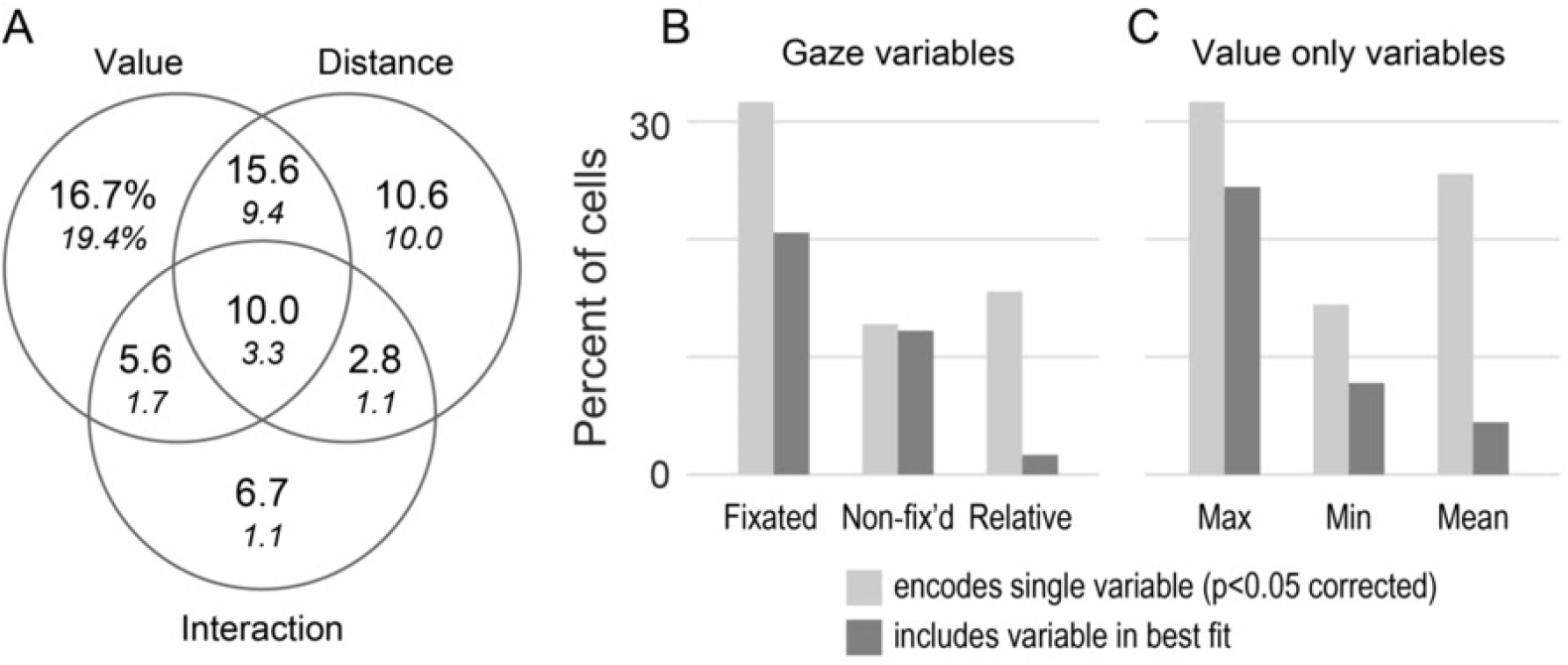
Modulation of OFC firing by gaze. (A) OFC activity in single-cue trials was explained by a linear model with three regressors: cue value, distance of fixation from the cue, and their interaction. The fraction of cells with significant effects are shown in the Venn diagram, using uncorrected and corrected thresholds of p<0.05 (large and small numerals, respectively). (B,C) OFC activity in two-cue trials was explained using a series of single- and two-variable models fit to every cell (Table 1). The light gray bars give the fraction of cells that encode each variable when fit by itself (out of n=180 cells, p<0.05 corrected). The dark gray bars give the fraction of cells that include a given variable in its best-fitting model.

### Neural recordings

Single electrodes were introduced into the brain through a sharpened guide tube whose tip was inserted 1-3mm below the dura. OFC was identified on the basis of gray/white matter transitions, and by consulting a high-resolution MRI acquired from each animal after chamber implantation. We targeted the fundus and lateral bank of the medial orbital sulcus and the laterally adjacent gyrus (Figure 5A), corresponding approximately to Walker’s area 13 (Öngür and Price, 2000).

From Monkey K, we recorded 116 neural unit signals ("cells”) over 25 sessions; and from Monkey F, we recorded 64 cells over 28 sessions. (Only sessions with concurrently collected neural data were used.) These included putative single units, characterized by large and well-isolated waveforms (n=63 from Monkey K, 44 from Monkey F), as well as multi-unit signals with low amplitude, poorly isolated waveforms (53 from K, 20 from F). Among the single units, some were isolated during the learning phase of the task (see above) and were selected for subsequent recording because they showed an apparent increase or decrease in firing during cue presentation; the remainder of the single units, and all multi-unit signals, were recorded without any prior observation of their activity during task performance. Neural data were only recorded and analyzed after the learning phase was complete. In some sessions several cells were recorded simultaneously by isolating more than one cell on a given electrode and/or by using two electrodes at once. See McGinty et al. (2016) for details.

After data collection, spikes were assigned offline to individual units based upon the principal component features of the waveforms (Plexon Offline Sorter 2.0). On rare occasions, cells initially designated as single units were re-categorized as multi-unit if they showed an abundance of short inter-spike intervals (more than 0.05% of intervals below 2ms). After unit sorting, the data were imported into MATLAB and the R software environment for analysis. There were no major differences in results obtain from single and multi-unit signals, and so their data are presented together.

### Analysis

#### Correlation between gaze allocation and reward anticipation

The objective of this analysis was to assess the trial-by-trial correlation between the fraction of gaze time devoted to Pavlovian cues and the fraction of time spent performing CRs in anticipation of reward delivery. Gaze and CR data were calculated in 500ms bins (50ms increments). Within each session, the across-trial correlation was calculated for all possible pairs of bins, and these correlations were then averaged across the sessions for each monkey (25 for Monkey K, and 28 for Monkey F). The resulting matrix of correlations has rows and columns correspond to the bin centers for gaze and CR data (respectively).

The correlation statistic was Spearman’s *rho*, a non-parametric, ranks-based measure of association that is outlier-resistant and suitable for non-normally distributed data. As a quality control measure, no correlation was calculated (i.e. the correlation was set to “nan”) when >80% of the gaze data or >80% of the CR data within a given bin had the same value; this happened most frequently in single “none” value trials, where the CR was often absent.

Correlation matrices are displayed as heatmaps showing the average correlation across sessions. One heatmap was calculated per single-cue trial type, and for each two-cue trial type one heatmap was calculated for the lower value cue and another for the higher value cue.

At each point on the map, the average correlation was compared to zero by means of a t-test. Red contours show points that surpass both an initial significance threshold of p<0.001 as well as a cluster-extent threshold of p<0.01 to control for multiple comparisons. The cluster extent threshold was determined as follows: For every map we created 1000 ‘null’ correlation maps using data in which the trial labels for the gaze data were randomly shuffled within each trial type. The null maps were thresholded at p<0.001, and the largest group of contiguous significant pixels (maximum cluster size) was recorded for each null map. (Contiguity was defined as a shared edge; shared corners were not considered contiguous.) This produced a distribution of 1000 maximum cluster sizes under the null hypothesis (no gaze-CR correlation). The cluster extent threshold was set to the 10th largest maximum cluster size (top 1.0 percentile). Then, in the original data, all clusters of significant points smaller than this threshold were discarded, corresponding to a cluster-level family-wise error rate (FWER) of p<0.01.

#### Gaze modulation of OFC neural activity

The objective of this analysis is to quantify the fraction of OFC neurons that are modulated shifts of gaze towards or away from the Pavlovian cues. Single cue trials were analyzed as a group, separately from two-cue trials.

##### Data source

The 180 neurons analyzed here are the subset of the 283 neurons analyzed in McGinty et al. (2016) for which both single- and two-cue data were collected. For single-cue trials, the main analyses of McGinty et al. 2016 are repeated here, and so the reported (Figure 5B,D and Figure 6A) are in essence a restatement of the findings of McGinty et al. 2016, in a subset of the original data. The two-cue trial data were obtained from these same 180 neurons, but have not been published before, with the exception of preliminary analyses in abstract form (McGinty et al., 2014).

##### Single cue trials

The basic unit of data was fixation-evoked firing, defined as the spike count observed 100-300ms after the beginning of each fixation epoch (see above). The temporal offset accounts for the typical delay in OFC responses to visual stimuli (Thorpe et al., 1983; Wallis and Miller, 2003).

For every neuron we fit the GLM in Equation 1. Cells with significant effects were identified for each regressor (p<0.05 both uncorrected and corrected with Holm’s Bonferroni). The GLM assumed a negative binomial error model, and is given by:

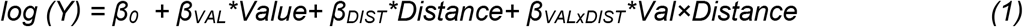

where each observation is a fixation (as defined above), *Y* is the gaze-evoked spike count for that fixation, *Value* refers to the volume of juice associated with the cue in each trial, *Distance* refers to the distance of gaze from the cue center for each fixation, and *Val* × *Distance* is the interaction of the Value and Distance variables (computed after centering them).

##### Two-cue trials

We focused on firing evoked by on-cue fixations (<=3 degrees from cue center), due to the low frequency of off-cue fixations (Figure 1G). To assess gaze and value effects, for every cell we fit GLMs that explained fixation-evoked firing on the basis of six variables (Table 1, columns). Three variables describe a value signal modulated by shifts of gaze between cues, i.e. a pattern of firing that depends not only on the values of the cues shown, but also on which cue is fixated at any given moment. They are: (#1) the value of the fixated target, (#2) the value of the other (non-fixated) target shown; and (#3) the relative value of the fixated target, defined as the fixated minus non-fixated target value, suggested by recent findings in frontal lobe recordings (Hunt et al., 2018). The three other variables describe a value signal with no modulation by gaze: (#4) the maximum of the two cue values shown, (#5) the minimum of the two shown, and (#6) the mean of the two shown.

**Table 1:** Models used to explain firing in two-cue trials. Each row specifies a model, which can use either one or two variables. The variables in a given model are indicated by ‘x’.

| Model | Gaze dependent variables |  |  | Non-gaze dependent variables |  |  | Number of cells best fit by each model |
| --- | --- | --- | --- | --- | --- | --- | --- |
|  | Fixated target | Non-fix'd target | Relative value | Maximum value | Minimum value | Mean value |  |
| 1 | x |  |  |  |  |  | 5 |
| 2 |  | x |  |  |  |  | 2 |
| 3 |  |  | x |  |  |  | 3 |
| 4 |  |  |  | x |  |  | 8 |
| 5 |  |  |  |  | x |  | 0 |
| 6 |  |  |  |  |  | x | 8 |
| 7 | x | x |  |  |  |  | 6 |
| 8 | x |  |  | x |  |  | 20 |
| 9 | x |  |  |  | x |  | 6 |
| 10 |  | x |  | x |  |  | 11 |
| 11 |  | x |  |  | x |  | 3 |
| 12 |  |  |  | x | x |  | 5 |

Because these six variables are not linearly independent, they cannot be assessed simultaneously in the same model. For example, the ‘mean value’ variable (#6 above) is a linear combination of the ‘max value’ and ‘min value’ variables (#4 and #5 above), meaning that independent estimates for these three variables cannot be obtained from a single model. We therefore adopted a competitive modelling approach, in which we fit a set of models containing either one or two regressors (see Table 1, rows 1-12), and the identified the best fitting model for each cell using Akaike’s information criterion (AIC). The results were quantified by finding the percentage of cells with significant effects of each single variable when fit by itself (light bars in Figure 6B,C), as well as the percentage of cells that included a given variable in its best-fit model (dark bars in Figure 6B,C).

The set of tested models is shown in Table 1. Other variable combinations were not tested due to the linear dependence of the variables. In brief: All models with >4 variables and some with 3 variables were excluded due to the strict linear dependence of the regressors, as in example above. The subset of 3-variable models that were able to be fit could not be distinguished from one another in terms of goodness-of-fit, because they all explained the same portion of variance in the data (again due to the non-independence of the regressors), and therefore yielded the same AIC. (In linear algebra terms, the matrix of regressors for fit-able 3-variable models all shared the same basis.) Some 2-variable models also yielded non-unique AICs for the same reason, and were also excluded. Thus, Table 1 shows all the combinations of the six regressors that can be fit in a single model and that also uniquely explain variance in firing -- and can therefore be compared in goodness-of-fit terms.

#### Correlation between OFC activity and reward anticipation

The objective of this analysis was to assess the trial-by-trial correlation between the activity of OFC neurons and fraction of time spent performing CRs in anticipation of reward delivery. This procedure is similar to the gaze-CR correlation calculation described above. Spiking and CR data were calculated in 500ms bins (50ms increments) and the correlation was performed across trials for all possible pairs of bins, yielding a matrix of correlation coefficients. No correlation was calculated (data set to “nan”) when >80% of the spike data or >80% of the CR data in a given bin had the same value. Correlations were calculated individually for each OFC cell (n=116 for Monkey K, 64 for Monkey F), and then averaged across cells. Six correlation matrices were calculated for each cell, one for each of the six trial types shown in Figure 1C and 1D.

Unlike the gaze-CR calculation, the correlation statistic was an unsigned variable that we term the “absolute adjusted correlation”, defined as:

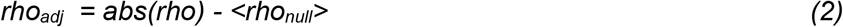

where abs() is the absolute value function, and *rho* is the raw Spearman’s correlation between the spike count and the CR. To find <rho_null_>, we randomly shuffle the trial labels 100 times within a cell, find the absolute Spearman’s correlation in each shuffle, and take the mean across shuffles. Thus, <rho_null_> is the absolute correlation that would be expected under the null hypothesis that spiking and the CR are unrelated; it is always above zero, because even totally random data produce spurious non-zero correlations. The rationale for using rhoadj is as follows: First, taking the absolute value of the raw spike-CR correlation puts positive spike-CR relationships (more spiking -> more CR) on the same scale as cells that have a negative spike-CR relationship (more spiking -> less CR); this allows us to average the correlations across all cells, regardless of the effect direction. Second, by subtracting <rho_null_>, the value of rho_adj_ is expected to be zero for cells in which there is no relationship between spiking and the CR, but is expected to be positive for cells in which there is a spike-CR relationship. The across-cell mean of rho_adj_ can be therefore assessed by a t-test versus zero, to determine whether a reliable spike-CR relationship exists at the population level.

The heatmap in Figure 7B shows the rho_adj_ for one trial type in one monkey, calculated at all pairs of time bins, averaged across all cells. The heatmap was thresholded at p<0.001, but there were no significant pixels that survived cluster correction to a family-wise error rate of p<0.01.

**Figure 7:**
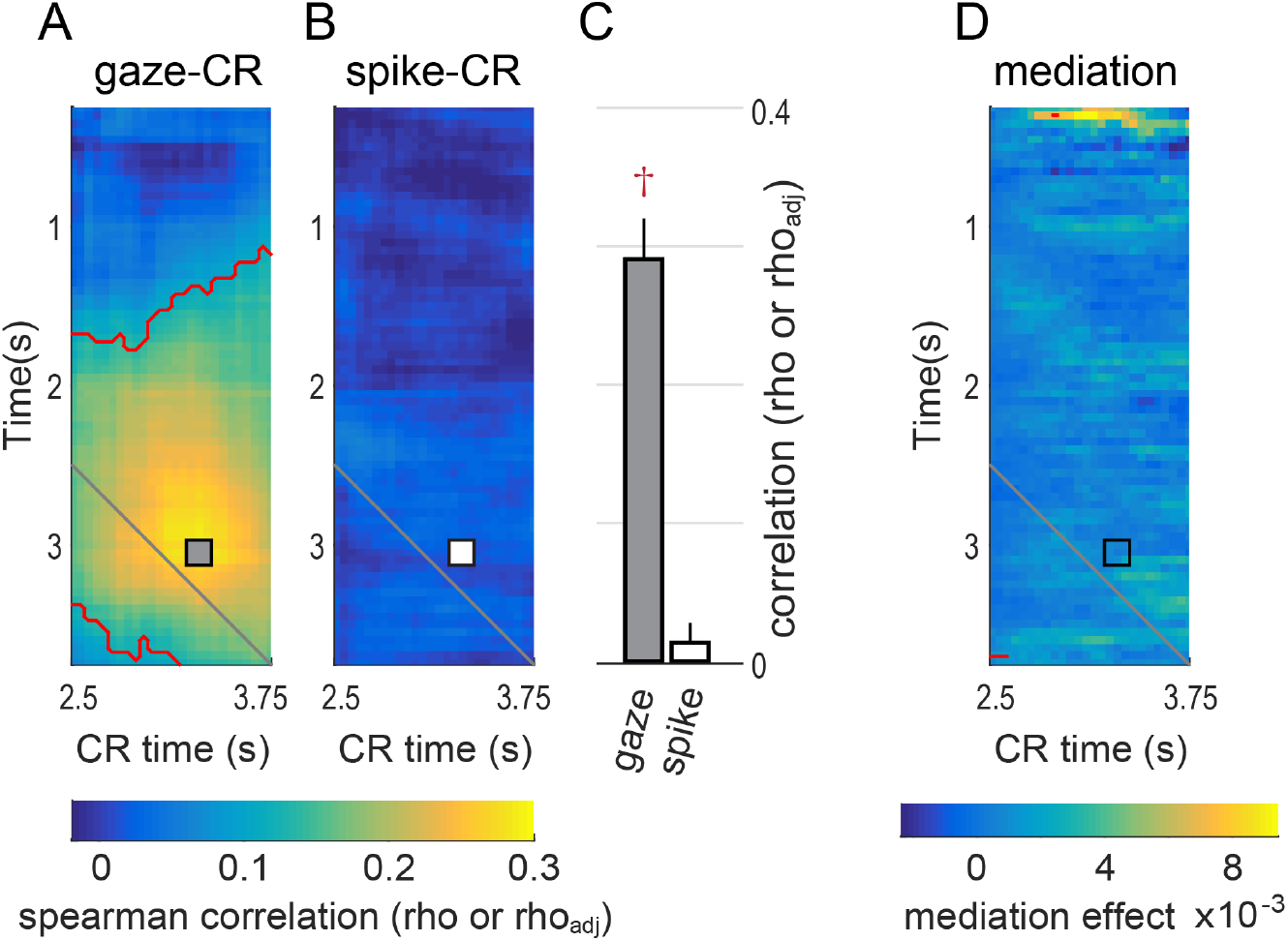
OFC activity does not predict CRs, and does not mediate the effects of gaze on CR, in example data from single ‘small’ cue trials in Monkey K. (A) The average gaze-CR correlation (rho) in single ‘small’ cue trials in Monkey K, reproduced from Figure 3, using the same conventions. The gray square shows the pixel with the highest correlation. This same pixel is marked with a square in panels B and C. Red contours indicate significant correlations as in Figure 3. (B) The cell-averaged spike-CR correlation (rho_adj_), using data from the same trial types and monkey as panel A. The white square corresponds to the peak gaze-CR effect in panel A.(C) The left and right bar give the mean correlations at the pixel marked with the square in panel A and B, respectively. Whiskers indicate SEM. These data are reproduced in Figure 8A, alongside data from all trial types in both monkeys. (D) Average mediation effects for single ‘small’ cues in Monkey K. See main text for details. Red contours indicate mean mediation effects above zero at p<0.001 (uncorrected) Mediation effects takes the same units as the regression estimates (beta values) upon which the analysis is based (see Methods).

In the bar graphs in Figure 8, the cell-averaged rhoadj is shown for all trial types and both monkeys, but at only a single point on the heatmap, selected as follows: For each of the gaze-CR heatmaps in Figure 4, we selected the point with the highest average correlation (black squares). In almost all trial types this peak point was above the diagonal, reflecting the fact that overt attention (gaze) tends to predict subsequent CRs. However, in the three single-cue trial types for Monkey F, there were gaze-CR effects of roughly equal magnitude both above and below the diagonal. For these conditions, we selected the peak within the above-diagonal data (black diamonds in Figure 4B), in order to maintain the temporal order of the predictive relationship that is the focus of the study (Figure 2). At these selected points we then computed the averaged spike-CR correlation across all cells, as in Figure 7B (*rho_adj_*), and compared it to zero by means of a t-test. Note that for every two-cue trial type, there are two gaze-CR matrices calculated (one for each cue, Figure 4), but only one spike-CR matrix. Therefore, to generate the data for two-cue trials in Figure 8, each spike-CR matrix is sampled at two points, one corresponding to the maximum gaze-CR effect for the lower value cue, and the other corresponding to the maximum effect for the higher value cue.

**Figure 8:**
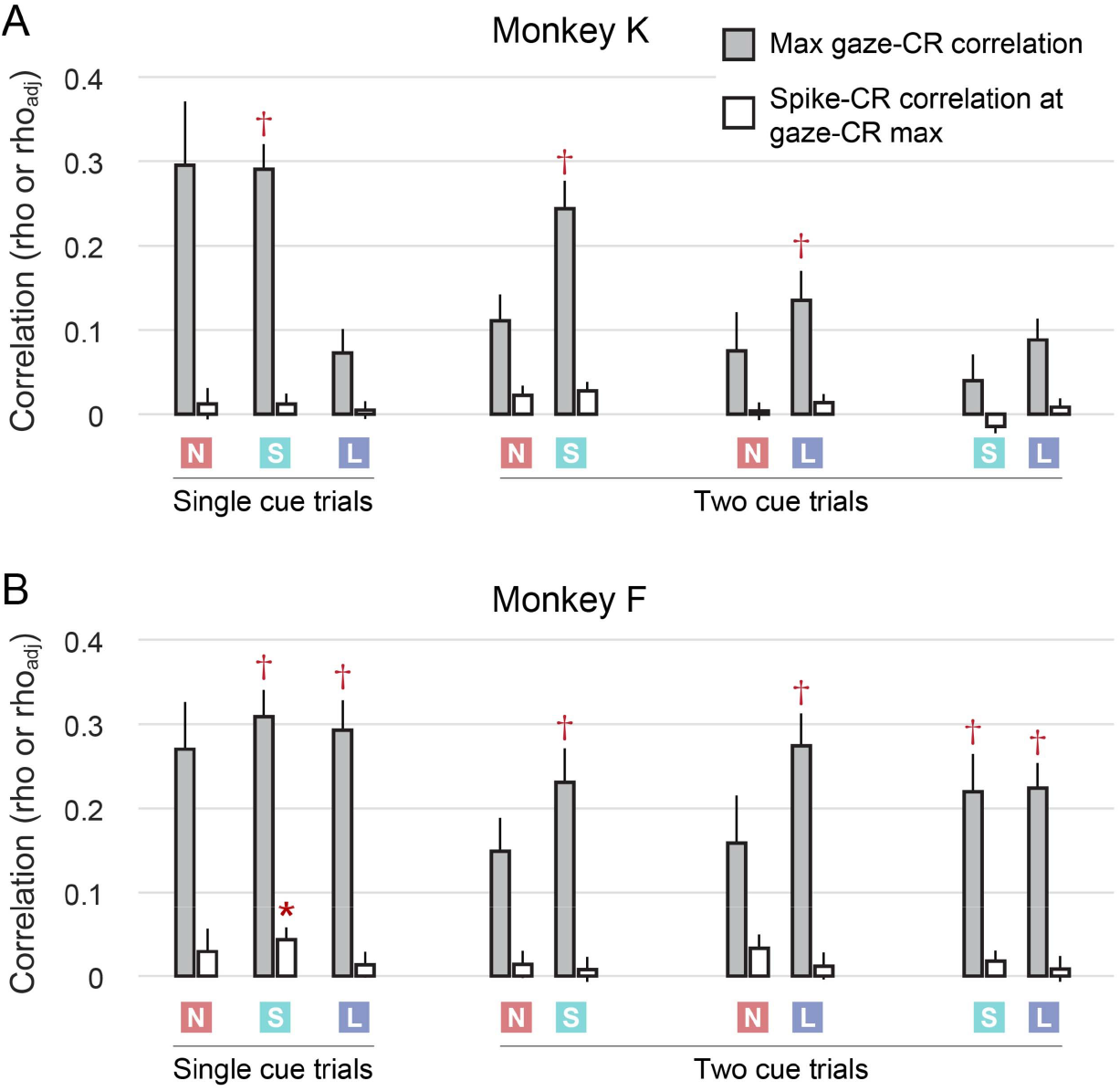
OFC neural activity is not correlated with CRs. Each pair of bars corresponds to one heatmap in Figure 4. The left bar gives the gaze-CR correlation at the time bins for which this effect was maximal (black squares in Figure 4); and the right bar gives the spike-CR correlation at these same time bins. The colored squares indicate whether the data pertain to ‘none’, ‘small’, or ‘large’ value cues (red N, green S, and blue L, respectively). For two-cue trial types, the effects are assessed separately for each cue, hence there are two sets of results for each two-cue trial type. The data from Figure 7C are reproduced in panel A (single ‘small’ cue trials). Whiskers give SEM. Daggers indicate that the peak gaze-CR effect falls within the significance contours in Figure 4. Asterisk indicates significant difference from zero, p<0.05, corrected.

#### Mediation analysis

The objective of this analysis was to quantify the degree to which OFC activity explains the correlation between gaze allocation and reward-anticipating CRs. The term ‘mediation’ is used in a purely statistical sense, and does not by itself imply a causal relationship between the variables. As with the gaze-CR analysis (Figure 4), a separate analysis was performed for each single-cue trial type, and two separate analyses were performed for each two-cue trial type (one each for the low and high value cue). Thus, nine separate mediation analyses were performed for each Monkey, corresponding to the nine gaze-CR analyses shown in Figure 4.

To quantify the mediation effect attributable to a single OFC cell, we measured gaze allocation to cues, spike counts, and CR data in 500ms bins. For every pair of time bins *x* and *y*, the following ordinary least squares linear models were fit:

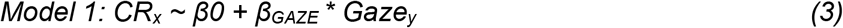

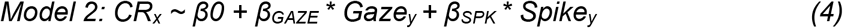

where *CR_x_* is the conditioned response observed within time bin *x*, *Gaze_y_* is the gaze allocation for a given cue in time bin *y*, and *Spike_y_* is the spike count from the cell in time bin *y* (observed concurrently with gaze). In this analysis gaze allocation and CR are quantified in units of time (range 0 to 500ms), such that the regression estimate β_GAZE_ can be interpreted as the linear effect of gaze on the CR. For example, a β_GAZE_ of 0.3 would indicate that for every 1 second increase in gaze allocation, an increase of 0.3 seconds in CR would be expected.

If β_GAZE_ is the same magnitude in both Model 2 and Model 1, this indicates that the *Spike* variable explains variance in the CR that is not attributable to the *Gaze* variable. In contrast, if βGAZE is smaller in Model 2 than in Model 1, it indicates that the the *Spike* variable has subsumed variance in the CR that would otherwise be accounted for by *Gaze*; this is evidence that *Spike* statistically mediates the linear association between the *Gaze* and *CR* variables. Thus, the mediation effect for a given cell at a given pair of time bins was calculated by subtracting the estimate βGAZE resulting from Model 2 from β_GAZE_ resulting from Model 1.

Nine mediation effect matrices were calculated for each OFC cell, and the median mediation effects across cells were compared to zero by means of a sign rank test. In Figure 7D, the heatmap shows the median matrix of mediation effects over n=116 cells, measured in single ‘small’ cue trials in Monkey K.

The mediation analysis was repeated in the subset of cells showing modulation by gaze. In this analysis we included any cells with significant effects (p<0.05 corrected) of fixation distance or the interaction variable in the analysis of single-cue trials; and any cells with significant effects (p<0.05 corrected) of one of the three fixation-related variables in the analysis of two-cue trials (Table 1, rows 1-3). A total of 84 cells were included, 45 from Monkey K and 39 from Monkey F.

## RESULTS

### Reward anticipation and gaze allocation to Pavlovian cues

Two monkeys performed the Pavlovian conditioning task in Figure 1. To begin a trial, monkeys briefly held their gaze on a fixation point, after which one or two visual cues appeared on the display for 4 seconds (Figure 1A,B). The monkeys were free to move their eyes throughout this period; eye movements were monitored, but had no effect on the trial outcome. At the end of 4 seconds, a juice reward was delivered as follows: single cues resulted in a guaranteed reward of 0, 1, or 3 drops (“none”, “small”, or “large”), with reward size determined by cue color (Figure 1C). Presentation of two different cues resulted in random delivery of one of the two indicated rewards, and the monkeys could not predict or influence which one would be delivered (Figure 1D). Single-cue and two-cue trials were randomly interleaved; cue selection was random, as was the placement of cues on the left and right sides of the display.

When the expected reward was non-zero, monkeys made Pavlovian conditioned licking responses (CRs) in anticipation of reward delivery, beginning approximately 2 seconds after the onset of the cue(s), and reaching a maximum just before reward delivery at 4 seconds (Figure 1E). On average, the CR magnitude increased monotonically with the mean value of the cue(s) shown (Figure 1F). Importantly, the average CRs differed significantly among the three single-cue trials, indicating that the monkeys successfully learned the individual cue-reward contingencies. We therefore consider these CRs to be indicators of the monkeys’ anticipation of reward on a given trial.

The allocation of the monkeys’ gaze -- where they looked and for how long -- was quantified by the fraction of time in every trial that the monkeys fixed their gaze on each cue. Gaze allocation varied as a function of cue value, but, unlike CRs, was not monotonically dependent on cue value (Figure 1G). Importantly, monkeys devoted non-zero fixation time onto ‘none’ cues, and also devoted non-zero fixation time onto the smaller of two cues shown simultaneously. Thus, the behavioral and neural effects of gaze allocation onto cues could be assessed regardless of cue value.

To summarize, monkeys were shown simple appetitive conditioned cues, either singly or in pairs. The monkeys allocated a significant portion of their gaze (overt attention) towards the cues, and performed anticipatory CRs commensurate with average cue value. The major questions of this study are whether trial-by-trial variability in gaze allocation corresponds to variability in CR magnitude (Figure 2A), and whether this correlation could be mediated by the value representations expressed in single OFC neurons (Figure 2B,C). In the next section, we document the trial-by-trial correlation between gaze allocation and CRs.

### Allocation of gaze to appetitive cues predicts trial-to-trial reward anticipation

On each trial, gaze allocation was defined as the fraction of time that the monkey spent looking at a cue (gaze < 3 degrees from cue center), and CRs were quantified according to the fraction of time that a licking response was detected. These two variables were measured in 500ms bins at 50ms increments during cue presentation, and the correlation between them was calculated across trials within a session for every possible pair of time bins in the trial. This yielded a measure of whether gaze allocation early in the trial was correlated with CRs later in the trial, and vice versa. Note that the CRs were only calculated at time bins with centers 2500ms postcue or later, due to the near total lack of CRs before this time (Figure 1E).

Correlations were calculated within a given session (n=28 for Monkey F, 25 for Monkey K), and then averaged across sessions. Because correlation patterns differed substantially according to cue value (see below), the correlations were calculated separately for each of the six trial types shown in Figure 1C,D (‘none’, ‘small’, ‘large’, ‘none-small’, ‘none-large’, ‘small-large’). Furthermore, in two-cue trial types, gaze allocation was tallied separately for each individual cue shown, permitting two separate correlation calculations to be obtained for every two-cue trial type. Therefore, for each monkey, nine total gaze-CR correlations were calculated: one for each trial type with a single cue, and two for each of the trial types with two cues.

#### Gaze-CR correlation in single trial type in a single subject

Figure 3A illustrates the trial-by-trial correlation between gaze allocation and Pavlovian CRs in one trial type (single ‘small’ cue) in Monkey K. The color of each pixel gives the session-wise mean correlation (*rho*) between the fraction of time the monkey spent looking at the cue, and the fraction of time a CR was detected; correlations were calculated across all of the trials within a given session, and then averaged across sessions (n=25). Points above the black solid diagonal line indicate gaze data that *precedes* (and could therefore predict) the CR data; and points below the black line indicate gaze data that *follows* CR data.

The highest average correlation occurred for gaze data measured at 3.05 seconds (y-axis) and CR data measured at 3.30 seconds (x-axis), with an average value of *rho*=0.291 (mean of 25 sessions, SEM 0.028). The positive correlation indicates that greater time spend gazing at the cue was associated with a larger CR; and the fact that the gaze data precedes the CR data indicates that gaze allocation could predict the upcoming CR on a trial by trial basis with ~0.25s temporal lag. In other words, the longer Monkey K looked at single ‘small’ cues at approximately 3 seconds in a trial, the more likely he was to exhibit an anticipatory CR 0.25 seconds later. Note that this predictive relationship was asymmetric: gaze was better able to predict CRs than the opposite, indicated by the higher correlations and greater fraction of significant pixels above the solid gray diagonal line compared to below it (Figure 3B).

#### All trial types and both subjects

The gaze-CR correlations for all subjects and trial types are shown in Figure 4, using the same conventions as Figure 3. The effects were highly variable from one condition to the next, with clear differences in the correlation patterns between subjects, and between conditions within each subject. However, despite the variability, two general patterns were evident. First, as in the example in Figure 3, the only significant correlations were positive (warm colors), meaning that a longer time spent looking at cues was associated with more frequent CRs. Significant positive correlations were found for 3 out of the 9 conditions in Monkey; and 7 out of 9 for Monkey F. No significant negative correlations were found. Second, gaze predicted CR performance to a greater degree than CRs predicted gaze behavior (above vs. below diagonal: 3/9 conditions in monkey K; 4/9 in monkey F). There were no conditions in which a significant difference was found in the opposite direction – i.e. in which CR predicted gaze to a greater extent than gaze predicted CR. In summary, we identified several trial conditions in which the longer that monkeys spent looking at (attending to) reward-associated cues, the greater their subsequent anticipation of reward delivery indicated by conditioned licking responses.

### Modulation of OFC neural activity by gaze

The positive correlation between overt attention to cues and Pavlovian CRs (Figures 3 and 4) must be explained by some neural mechanism that links these two behaviors. To identify this mechanism requires finding, at a minimum, neural activity related to the two behaviors of interest (Figure 2B,C). Here, we show that OFC firing is modulated by shifts of gaze towards or away from Pavlovian cues. Because the gaze-CR correlation was present in both single- and two-cue trials, it was necessary to test the effects of gaze on OFC activity in both kinds of trials. Single-cue gaze effects were shown in a prior study, and are recapitulated here using a subset of the original data (McGinty et al., 2016), consisting of those cells in which both single- and two-cue trials were tested. Data from two-cue trials are shown for the first time, and are analyzed separately from single-cue trials due to differences in gaze behavior when two cues are shown rather than one (see below). A total of 116 neurons were recorded in Monkey K, and 64 in Monkey F.

#### Single cue trials

In single-cue trials, monkeys typically shifted their gaze many times during the 4-second cue presentations, fixating at various locations on the display, both on and off the cue (Figure 1G,F; see also McGinty et al. Figure 2). To assess neural activity with reference to gaze location, OFC firing was measured from 100-300ms after the onset of each fixation; this ‘fixation-evoked’ firing was the basic unit of data for this analysis.

We fit a GLM that explained firing as a linear function of the value of the cue shown, the distance of gaze from the cue, and the value-by-distance interaction (Equation 1). The single cells in Figure 5B and 5D illustrate the encoding of all three variables: firing was greatest for fixations near to the cue (distance encoding), and was monotonically related to the volume of juice reward (value encoding). Critically, the effect of value was greatest for near-to-cue fixations, which constitutes an interaction between the value and distance effects. At the population level, large portions of neurons were significantly modulated by cue value (47.8 and 33.9% with GLM effects at p<0.05, uncorrected and corrected, respectively) and fixation distance from the cue (38.9 and 23.9%), and a smaller portion were modulated by the interaction term (25.0 and 7.2%). The mean regression estimate for the distance effect was negative (−0.0096, SEM 0.0020, p = 2*10^-6^), indicating that near-cue fixations elicited greater overall firing than fixations away; and the value estimate was not significantly different from zero (mean 0.032 SEM 0.050, p=0.62), indicating that neurons were equally likely to increase firing with cue value (as in Figure 5B) as they were to decrease firing (Figure 5D). Many neurons had more than one significant effect, indicated in the Venn diagram in Figure 6A (compare to McGinty et al. Figure 5A).

#### Two-cue trials

In two-cue trials, monkeys shifted their gaze throughout cue presentation. However, unlike in single cue trials, the majority of fixations were directed onto the cues (Figure 1G), leaving too few off-cue fixations to assess the effects of gaze distance. The analysis therefore used only firing evoked by on-cue fixations, again within a 100-300ms window after each. On-cue fixations are parameterized by the values of the fixated and non-fixated cue, so that gaze effects can be assessed by the degree to which firing is modulated by either of these variables. In addition, we assessed modulation by cue value in a non-gaze-dependent manner, given that during single-cue trials we found cells that encode only cue value with no effect of gaze distance (Fig 6A, McGinty et al. 2016 Figure 5J).

To identify gaze-modulated cells in two-cue trials, we began by fitting three GLMs per cell, each with a single variable that explained firing on the basis of cue values and the location of fixation. The first GLM explained firing according to the value of the cue targeted in a given fixation (‘fixated value’). Two examples of cells with significant (p<0.05 corrected) effects in the fixated value GLM are shown in Figure 5. The cell in Figure 5C fires more for fixations onto the higher of the two cues shown, whereas the cell in Figure 5E fires more for the lower of the two. The second GLM explained firing as a function of the value of the non-fixated target (‘nonfixated value’); and the third explained firing as a function of the value difference between the fixated and non-fixated target (‘relative value’, see Methods). In all 36.7% (n=66) OFC cells showed significant effects of at least one of the three GLMs described above, with the large majority of these showing significant modulation by fixated value (Figure 6B). Therefore, just as in single-cue trials, a substantial portion of OFC neurons have value signals that are modulated by gaze when two cues are shown.

While the primary objective of this analysis was to identify gaze-modulated cells in two-cue trials, additional analyses below provide a more comprehensive view of the variables encoded in this phase of the task. Specifically, we assess: encoding of non-gaze-dependent value variables; the mixture of variables encoded in single neurons; and the consistency of encoding between the single- and two-cue phases of the task.

First, to identify OFC cells modulated only by value (with no effect of gaze) we fit three additional GLMs with single variables that depended on the value of the visible cues, but not on which cue was targeted by a given fixation. These were: the maximum value of the two cues; the minimum of the two; and the mean of the two. In all, 33.9% (n=61) of OFC cells had significant effects (p<0.05 corrected) of at least one of these three ‘value only’ variables (Figure 6C).

In the single cue analysis, the variables of interest were mixed at the single-cell level, i.e. many single neurons were modulated by more than one variable. In the two-cue data, it was not possible to fit all six variables in the same GLM due to linear dependency among the regressors (see Methods). Therefore, to quantify mixed encoding of gaze-dependent and value-only variables, we used a competitive modelling approach. First we identified all cells that had a significant effect in at least one of the six single-variable GLMs above (n=77, p<0.05 corrected). Then, in these cells we also fit a set of two-variable GLMs using different combinations of the six variables (Table 1, rows 7-12). Finally, we calculated the goodness of fit (AIC) for all single-variable and two-variable models, and identified which model provided the best fit for a given cell. The results are shown in Table 1, and Figure 6B,C. The model that provided the best fit for the most cells was the two-variable model including ‘fixated value’ and ‘maximum value’ (n=20, 11% of all cells), which is consistent with these two variables producing the most significant effects when fit individually (Figure 6B,C). In all, 40 cells (22%) were best fit by a two-variable model that included one gaze-related variable and one value-only variable. Thus, as in single-cue trials, OFC neurons encode a mixture of task variables when two cues are shown.

Finally, we found that the same cells tended to be modulated in both the single- and two-cue task phases: 83 cells were modulated by at least one of the three single-cue analysis variables (p<0.05 corrected), 77 were modulated by at least one of the six two-cue analysis variables, and 53 were modulated by at least one variable in each task phase, which was significantly greater than expected by chance (35.5 expected, p=10^-7^ by chi-squared test). This was also true when considering neurons with gaze modulation: 48 cells were significantly modulated by either fixation distance or the distance-by-value interaction in the single-cue analysis, 66 were modulated by at least one of the three gaze-dependent variables in the two-cue analyses, and 30 were gaze modulated in both task phases, significantly more than expected by chance (17.6, p=10^-5^). Finally, the sign of value modulation was consistent across the population: assessed in all 180 cells, the regression estimates for value obtained in the single cue analysis was highly correlated with the regression estimates for ‘fixated value’ (r = 0.55), ‘maximum value’ (r=0.72), and ‘mean value’ (0.69) obtained in the two-cue analysis.

#### Summary

Shifts of gaze during the presentation of Pavlovian conditioned cues influenced the firing OFC neurons in a 100-300ms window following the onset of each fixation. When single cues were presented, many OFC cells encoded the distance of gaze from the cue, or expressed value signals that were modulated according to gaze distance. When two cues were presented, a portion of OFC cells encoded either the value of the cue fixated at any given moment, the value of the other (non-fixated) cue, or both of these variables (relative value).

### OFC neural activity does not predict reward anticipation

Our central hypothesis is that shifts of gaze influence Pavlovian CRs through the modulation of OFC neural activity (Figure 2). Above, we establish the first part of this mediation relationship (gaze shifts modulate OFC activity, Figure 5, 6). Here, we test the second arm of the mediation relationship, between OFC neural activity and CRs. To preview the results in brief: we find that on average OFC activity is only weakly predictive of CRs, and appears insufficient to act in a mediating role.

As in the gaze-CR analysis, we measured CRs (fraction of time a licking response was detected) and OFC activity from individual cells (spike count) in 500ms bins. We then calculated the across-trial correlation between spike count and CRs, using an unsigned correlation metric (rho_adj_) that takes a positive value regardless of whether spiking increases or decreases with respect to the CR, thereby placing all OFC cells on the same scale (see Methods). This was done for every possible pair of time bins, yielding a matrix of spike-CR correlations across different time points in the trial. A separate spike-CR correlation matrix was created for each of the six trial types show in Figure 1C and 1D. These were then averaged across all recorded cells.

The heatmap Figure 7B shows the spike-CR correlation matrix for single ‘small’ reward trials in Monkey K, averaged across 116 cells. Unlike the gaze-CR correlation for these trials (reproduced in Figure 7A), the spike-CR correlations are not statistically different from zero, and show no overall temporal pattern. In other words, within this example data, gaze explains variance in CRs, but concurrently observed neural activity does not. Intuitively, the weak correlations in Figure 7B and mismatch with respect to Figure 7A are evidence against a mediation relationship. We performed two analyses to quantify this intuition. First, we identified the time bins with the strongest gaze-CR correlation, and then asked whether the spike-CR correlation at this point was significantly above zero. In Figure 7A, this point is marked with a gray square, at x=3.30s and y=3.05s. The gaze-CR correlation at this point is *rho*=0.291 (mean of 25 sessions, SEM 0.028, significantly greater than zero by t-test, p<10-9). In contrast, the spike-CR correlation at this point is only *rho_adj_* = 0.012 (n=111 OFC cells, SEM 0.011), and is not significantly above zero (p = 0.26 by t-test). In other words, even when the predictive effect of gaze was strongest, there was no corresponding predictive effect in the OFC spiking data. We repeated this analysis in all trial types for both monkeys, selecting the time points at which the predictive correlation of gaze for CRs was maximal (see Methods). As shown in Figure 8, none of the spike-CR correlations at the selected points were significantly above zero in Monkey K, and only one point was significantly above zero for Monkey F (single ‘small’ cue trials, rho_adj_ = 0.043, n=55 OFC cells, SEM 0.014, p=0.042, corrected for multiple comparisons). Thus, there was virtually no evidence that OFC mediated the predictive relationship between gaze and CRs, even when this effect was maximal within a given trial type.

To confirm this result, we directly quantified the mediating effect of OFC with a mediation model, and expanded the scope of the analysis to consider all time bins (not just the gaze-CR maximum). For the example data (‘small’ trials for monkey K), the results are shown in the heatmap in Figure 7D. Positive values indicate evidence in favor of a mediation relationship (see Methods). The mediation effect at the maximum gaze-CR point (black square) was not significantly above zero (median 2.7×10^-4^, SEM 2.8*10^-3^, p = 0.30, n=114 cells). We repeated this analysis in all trial types in both monkeys, and found similar weak effects (not shown), with the strongest effect occurring in single ‘small’ cue trials in Monkey F (median 0.0064, SEM 0.0049, n=54 cells, p=0.043 uncorrected, p=0.78 corrected).

Moreover, mediation effects were weak at virtually all time points. In the example heatmap (Figure 7D) very few points show significant effects surpassing even an uncorrected threshold of p<0.001 (i.e. before cluster correction as was done in Figure 3 and 4), and most of these lie well below the diagonal and are therefore inconsistent with the mediation hypothesis, which dictates that gaze (and spiking) temporally precede CR behavior (Figure 2). The heatmap of mediation effects in Figure 7D was representative of the results from other trial types in both monkeys (not shown), that is, mediation effects were weak overall, and only a tiny fraction of points showed effects that were significantly above zero (p<0.001).

Two additional confirmatory analyses were performed. First, our central hypothesis (Figure 2) holds that mediation effects should be evident only in cells that are modulated by gaze, which we identify in Figure 6A and 6B. However, even when we averaged the mediation effects of only gaze-modulated cells (n=45 for monkey K and n=39 for monkey F, see Methods), the results were the same: no significant mediation at the points with the strongest gaze-CR effects, as well as weak mediation effects overall (not shown).

In summary, while we found strong evidence that shifts of gaze towards Pavlovian cues positively predicted reward-anticipating CRs, we found no evidence that CRs could be predicted by concurrently observed OFC firing, and, by extension, no evidence that OFC firing participates in the neural mechanisms that links attention and CRs.

## DISCUSSION

Neurons in the primate OFC represent the value of appetitive and aversive Pavlovian conditioned stimuli, and can predict the performance of a Pavlovian conditioned response (Morrison and Salzman, 2009). This suggests a role for OFC in the subjective value signals that ultimately inform Pavlovian responding. Our prior work showed that OFC value signals were modulated by moment-to-moment shifts of gaze (overt attention) towards Pavlovian cues, but left open the question of how this attentional modulation ultimately influences behavior. This was the core question of the current study, a timely issue in light of the clear effects of attentional shifts in economic decisions (Krajbich et al., 2010; Towal et al., 2013; Vaidya and Fellows, 2015), another form of appetitive motivated behavior. Our results therefore inform the larger effort to untangle the complex relationship between attention, neural value signals, and behavior.

In monkeys performing an appetitive Pavlovian conditioning task, gaze allocation positively predicted CRs: the longer the monkeys spent looking at a conditioned cue, the greater the likelihood that they would perform a conditioned licking response later in the trial -- though this effect differed between monkeys, and between trial types within monkey (Figure 4). OFC neural activity in this task was modulated by shifts of gaze, both for single cues presented alone (as reported earlier (McGinty et al., 2016)), and for cues presented in pairs. However, OFC activity did not predict conditioned licking responses on a trial-by-trial basis, and as a result we found no evidence that OFC firing could mediate the effects of attention on conditioned responses. Below we discuss each of these findings in depth.

### Attention predicts conditioned responses performed in anticipation of reward

When it was present, the correlation between gaze and CR was always positive, meaning that more gaze devoted to a cue was associated with more frequent CRs. In addition, gaze *predicted* subsequent CRs to a greater extent than the opposite. Together, these suggest that gaze enhances the subjective value of Pavlovian cues, similar to the effects on objects offered during economic choice. However, this conclusion comes with several important caveats. First, we did not directly manipulate gaze, and so cannot conclude with certainty that it has a causal effect on CRs. Second, the correlations were not uniform in both subjects: Whereas Monkey F had consistent effects for both ‘small’ and ‘large’ value cues, effects in Monkey K were prominent for ‘small’ cues, and weak or absent for ‘large’. In addition, for single cue trials in Monkey F, CR data appeared to predict gaze behavior to a similar extent that gaze predicted CRs. However, despite these differences, two key patterns are present in both subjects: the positive sign of the correlation, and the overall greater predictive effect of gaze for subsequent CRs.

The source of variability in the gaze-CR correlations is not immediately clear, given the inconsistency between the two subjects, in particular, the negligible effects for ‘large’ value cues in Monkey K, in comparison to those in Monkey F. One possibility is that for Monkey K, the largest available reward amounts to a form of “jackpot” for which the subjective value is maximal and essentially inelastic. This is consistent with observations in humans that larger rewards are subject to shallower discount functions than smaller rewards (Green et al., 1999).

The inconsistency of gaze-CR effects also makes it difficult to specify the nature of the neural mechanism that links attention and reward anticipation. In economic choice, computational models support a multiplicative mechanism in which gaze effects increase as a function of the value of the attended item (Krajbich et al., 2010; Towal et al., 2013; Vaidya and Fellows, 2015). The data of Monkey K are inconsistent with a multiplicative mechanism, with virtually no gaze effects for the highest value cue. Our results also appear to rule out a simple additive mechanism (Cavanagh et al., 2014), because the overall weak effects on ‘none’ value cues suggests that gaze must interact with, rather than simply amplify, neural value representations that underlie CRs. Additionally, gaze effects in both monkeys appear to be sensitive to whether cues are presented alone or in pairs: Monkey K shows a positive gaze-CR effect for ‘small’ cues presented alone, but not when presented alongside ‘large’ value cues; and Monkey F shows a positive gaze-CR effect for ‘none’ cues presented alone, but not alongside another cue. This suggests that gaze effects may depend on the relative cue value.

In summary, attending to Pavlovian cues appears to enhance their subjective value. While this broadly similar to attentional effects in economic choice, the underlying mechanisms may differ. Additional experiments using a greater variety of reward values, and a larger number of subjects, may be necessary to resolve this question.

### Attention modulates OFC value signals

OFC value signals for single cues are modulated by overt shifts of attention towards or away from those cues (McGinty et al., 2016). The current study extends these findings to gaze shifts between two items of different value. In general, the effects of gaze were consistent in both single- and two-cue contexts. In both contexts, many OFC cells represented cue value in some form, but only a subset were also modulated by shifts of gaze. In two-cue trials, attended items were preferentially represented, indicated by the fact that the ‘fixated value’ variable was represented by greater fraction of cells than ‘non-fixated value’ (Figure 6B). This is consistent with gaze effects in single cue trials, where OFC neurons express a stronger distinction between the cue values when the monkeys gaze towards the cues, illustrated in the example cells in this study (Fig 5B,D), and in cell-averaged data in McGinty et al. Figure 5D-E. The preferential representation of attended over unattended items is also consistent with the effects of covert shifts of attention, obtained under similar Pavlovian-like conditions (Xie et al., 2018). Our findings are also consistent with those of Hunt et al. (2018), who report OFC value signals that primarily reflect the fixated object during a decision-making task. In contrast, one recent report (Rich and Wallis, 2016) shows no effect of fixation on OFC value signals; this may be attributable to the fact that, unlike in Hunt et al. 2018, fixation onto the choice options was not a requirement for successful task performance.

In summary, whether one or two value-associated objects are present, a substantial portion of the OFC’s value representation is modulated by shifts of gaze during natural free viewing. An open question is whether the effects of overt attention and those of covert attention are two facets of the same common neural mechanism, or result from two distinct neural mechanisms under different experimental contexts -- i.e. free viewing versus enforced fixation.

### Potential mechanisms for attentional modulation of conditioned responses

OFC activity was almost entirely uncorrelated with performance of CRs, meaning there is no evidence that the OFC mediates the predictive relations between gaze and CRs. This negative finding is at odds with the observations of Morrison and Salzman (2009), who found that responses of OFC neurons to Pavlovian cues could indeed predict the performance of conditioned responses on a trial-by-trial basis. The discrepancy may be explained by a difference in task design: First, in Morrison and Salzman the visual cues appeared only briefly (~300ms), followed by a trace interval of 1.5 seconds with no stimuli present. Second, both appetitive and aversive unconditioned stimuli were used (juice reward and an air puff), corresponding to two distinct CRs. Thus, to perform the task optimally, the monkeys were required to remember the conditioned cue over the trace interval, and to perform the correct response in anticipation of the associated outcome. In contrast, our task had no mnemonic requirement, and only one possible outcome. It is possible that the greater working memory and behavioral demands of the task used by Morrison and Salzman may have required greater recruitment of prefrontal circuitry, and therefore a produced measurable correlation between OFC activity and behavior.

Negative findings are not in themselves evidence of no effect. Two follow-up experiments could potentially clarify these findings. First, simultaneous recording from multiple neurons would allow for less noisy estimates of subjective value signals on single trials, and therefore less noisy estimates of the mediation effects. This approach may be particularly appropriate for OFC, where individual cells appear to be poor estimators of underlying value variables due to high within-cell noise and low across-cell correlation (Conen and Padoa-Schioppa, 2015). Second, direct manipulation of OFC neurons would establish whether gaze-CR predictions depend on normal OFC function. Interestingly, there is preliminary evidence that Pavlovian-conditioned pupillary responses in primates are attenuated by OFC lesions, suggesting involvement for OFC in multiple forms of conditioned responding (Hwang et al., 2018).

If the negative mediation findings are indeed reliable, then regions other than OFC must form the neural mechanism linking gaze and reward anticipation. These may be regions that are both involved in consumatory oromotor movements (i.e. the licking response measured in this study), and are also subject to attentional modulation. One candidate region is the ventral striatum (VS). Although oromotor responses to pleasant and unpleasant tastes do not require the VS (or any circuitry above the midbrain (Grill and Norgren, 1978)), they can be evoked or repressed by pharmacological manipulation of the VS, especially of dopamine and opioid receptors in the rostral shell of the nucleus accumbens (Castro et al., 2015). VS manipulations also modulate the incentive value of Pavlovian conditioned stimuli (Corbit et al., 2001), and influence reward-seeking behavior in response to those stimuli (Nicola, 2010; Hoffmann and Nicola, 2014). In a human imaging study by Lim et al. (2011), value representations evident in the VS BOLD signal were modulated by gaze shifts between visual objects of differing value, similar to the modulation exhibited by OFC neurons in this study. Thus, the VS exhibits both attention-modulated value signals and exerts top-down control over appetitive oromotor responses, making it a candidate region for mediating the effects of attention on reward anticipation we observed in this study.

Other candidates are regions projecting to the VS, particularly the ventromedial prefrontal cortex (vmPFC), which projects to the shell of the nucleus accumbens (Heilbronner et al., 2016) and also express gaze-modulated value signals (Lim et al., 2011). In contrast, the region recorded in this study, Walker’s area 13, projects primarily to the ventromedial caudate nucleus and accumbens core, with only weak projections to the accumbens shell (Haber et al., 1995). Area 13’s direct connections with the vmPFC are also sparse (Carmichael and Price, 1996; Öngür and Price, 2000). Thus, the overall limited connectivity between the recorded region and putative oromotor output centers may account for the very weak correlations we observed between neural activity activity and licking CRs. By this logic, we predict that neurons in vmPFC should exhibit stronger correlations with licking CRs than neurons in OFC.

### Implications for decision-making and other motivated behaviors

Overt shifts of attention influence another important form of motivated behavior: economic choice. During decision-making, choice is biased in favor of the item fixated (attended to) first in a given trial, as well as towards the item that received the largest overall portion of the total fixation time prior to the choice (Krajbich et al., 2010; Towal et al., 2013; Vaidya and Fellows, 2015; Tavares et al., 2017). This effect is well-explained by serial sampling models in which fixation biases the accumulation of evidence in favor of whichever item is attended at any given moment (Krajbich et al., 2010; Tavares et al., 2017). Neurons that preferentially encode the value of attended stimuli -- like those reported here -- would be an important element of such a mechanism, providing input to downstream circuitry (presumably proximal to motor outputs) in which the evidence accumulation takes place. In theory, a similar mechanism could underlie gaze effects on Pavlovian responses. However, our results suggest that the OFC value signals do not perform this function in Pavlovian contexts.

These results therefore suggest two related questions that must be addressed in future work: The first concerns the precise locus of attention-modulated value signals that ultimately influence behavior. The OFC appears to not be involved in gaze modulation of Pavlovian responses, but its role in gaze effects on economic choice is unclear. And the second concerns the extent to which the attentional effects on choice and Pavlovian responses are attributable to common neural substrates. As we note above, the inconsistent gaze modulation for ‘large’ value cues would not be expected according to computational models of choice, suggesting at least one major mechanistic difference between choice and Pavlovian contexts.

## ACKNOWLEDGEMENTS

The author wishes to thank W.T. Newsome for funding, material support, and helpful discussion in preparation of the manuscript J. Brown, S. Fong, J. Powell, J. Sanders, and E. Carson provided technical assistance. We thank D. Moorman, and D. Kimmel for helpful comments.

## REFERENCES

Carmichael ST, Price JL (1996) Connectional networks within the orbital and medial prefrontal cortex of macaque monkeys. J Comp Neurol 371:179–207.

Castro DC, Cole SL, Berridge KC (2015) Lateral hypothalamus, nucleus accumbens, and ventral pallidum roles in eating and hunger: interactions between homeostatic and reward circuitry. Front Syst Neurosci 9 Available at: https://www.frontiersin.org/articles/10.3389/fnsys.2015.00090/full [Accessed April 23, 2018].

Cavanagh JF, Wiecki TV, Kochar A, Frank MJ (2014) Eye Tracking and Pupillometry are Indicators of Dissociable Latent Decision Processes. J Exp Psychol Gen 143:1476–1488.

Conen KE, Padoa-Schioppa C (2015) Neuronal variability in orbitofrontal cortex during economic decisions. J Neurophysiol 114:1367–1381.

Corbit LH, Muir JL, Balleine BW (2001) The Role of the Nucleus Accumbens in Instrumental Conditioning: Evidence of a Functional Dissociation between Accumbens Core and Shell. J Neurosci 21:3251–3260.

Engbert R, Kliegl R (2003) Microsaccades uncover the orientation of covert attention. Vision Res 43:1035–1045.

Gidlöf K, Anikin A, Lingonblad M, Wallin A (2017) Looking is buying. How visual attention and choice are affected by consumer preferences and properties of the supermarket shelf. Appetite 116:29–38.

Green L, Myerson J, Ostaszewski P (1999) Amount of reward has opposite effects on the discounting of delayed and probabilistic outcomes. J Exp Psychol Learn Mem Cogn 25:418–427.

Grill HJ, Norgren R (1978) The taste reactivity test. II. Mimetic responses to gustatory stimuli in chronic thalamic and chronic decerebrate rats. Brain Res 143:281–297.

Haber SN, Kunishio K, Mizobuchi M, Lynd-Balta E (1995) The orbital and medial prefrontal circuit through the primate basal ganglia. J Neurosci 15:4851–4867.

Heilbronner SR, Rodriguez-Romaguera J, Quirk GJ, Groenewegen HJ, Haber SN (2016) Circuit Based Cortico-Striatal Homologies between Rat and Primate. Biol Psychiatry Available at: http://www.sciencedirect.com/science/article/pii/S0006322316323885 [Accessed June 26, 2016].

Hoffmann J du, Nicola SM (2014) Dopamine Invigorates Reward Seeking by Promoting Cue-Evoked Excitation in the Nucleus Accumbens. J Neurosci 34:14349–14364.

Hunt LT, Malalasekera WMN, Berker AO de, Miranda B, Farmer SF, Behrens TEJ, Kennerley SW (2018) Triple dissociation of attention and decision computations across prefrontal cortex. Nat Neurosci 21:1471–1481.

Hwang J, Noble P, Murray E (2018) Orbitofrontal cortex lesions disrupt anticipatory autonomic responses to reward magnitude in macaque monkeys. In. San Diego, CA: Society for Neuroscience.

Kimmel DL, Mammo D, Newsome WT (2012) Tracking the eye non-invasively: simultaneous comparison of the scleral search coil and optical tracking techniques in the macaque monkey. Front Behav Neurosci 6:49.

Krajbich I, Armel C, Rangel A (2010) Visual fixations and the computation and comparison of value in simple choice. Nat Neurosci 13:1292–1298.

Lim S-L, O’Doherty JP, Rangel A (2011) The Decision Value Computations in the vmPFC and Striatum Use a Relative Value Code That is Guided by Visual Attention. J Neurosci 31:13214–13223.

McGinty V, Rangel A, Newsome W (2014) Do orbitofrontal cortex neurons encode relative value during free gaze? In. Washington, DC.

McGinty VB, Rangel A, Newsome WT (2016) Orbitofrontal Cortex Value Signals Depend on Fixation Location during Free Viewing. Neuron 90:1299–1311.

Morrison SE, Salzman CD (2009) The Convergence of Information about Rewarding and Aversive Stimuli in Single Neurons. J Neurosci 29:11471–11483.

Nicola SM (2010) The Flexible Approach Hypothesis: Unification of Effort and Cue-Responding Hypotheses for the Role of Nucleus Accumbens Dopamine in the Activation of Reward-Seeking Behavior. J Neurosci 30:16585–16600.

Öngür D, Price JL (2000) The Organization of Networks within the Orbital and Medial Prefrontal Cortex of Rats, Monkeys and Humans. Cereb Cortex 10:206–219.

Rich EL, Wallis JD (2016) Decoding subjective decisions from orbitofrontal cortex. Nat Neurosci advance online publication Available at: http://www.nature.com/neuro/journal/vaop/ncurrent/full/nn.4320.html [Accessed June 10, 2016].

Tavares G, Perona P, Rangel A (2017) The Attentional Drift Diffusion Model of Simple Perceptual Decision-Making. Front Neurosci 11 Available at: http://journal.frontiersin.org/article/10.3389/fnins.2017.00468/full [Accessed September 20, 2017].

Thorpe SJ, Rolls DET, Maddison S (1983) The orbitofrontal cortex: Neuronal activity in the behaving monkey. Exp Brain Res 49:93–115.

Towal RB, Mormann M, Koch C (2013) Simultaneous modeling of visual saliency and value computation improves predictions of economic choice. Proc Natl Acad Sci 110:E3858–E3867.

Vaidya AR, Fellows LK (2015) Testing necessary regional frontal contributions to value assessment and fixation-based updating. Nat Commun 6:10120.

Wallis JD, Miller EK (2003) Neuronal activity in primate dorsolateral and orbital prefrontal cortex during performance of a reward preference task. Eur J Neurosci 18:2069–2081.

Xie Y, Nie C, Yang T (2018) Covert shift of attention modulates the value encoding in the orbitofrontal cortex. eLife 7:e31507.

